# Improved computational epitope profiling using structural models identifies a broader diversity of antibodies that bind the same epitope

**DOI:** 10.1101/2023.06.09.543890

**Authors:** Fabian C. Spoendlin, Brennan Abanades, Matthew I. J. Raybould, Wing Ki Wong, Guy Georges, Charlotte M. Deane

## Abstract

The function of an antibody is intrinsically linked to which epitope it engages. Clonal clustering methods, based on sequence identity, are commonly used to group antibodies that will bind the same epitope. However, such methods neglect the fact that antibodies with highly diverse sequences can exhibit similar binding site geometries and engage common epitopes. In a previous study we described SPACE1, a method that structurally clustered antibodies in order to predict their epitopes. This methodology was limited by the inaccuracies and incomplete coverage of template-based modelling. It was also only benchmarked at the level of domain-consistency on one virus class. Here, we present SPACE2, which uses the latest machine learning based structure prediction technology combined with a novel clustering protocol and benchmark it on binding data that has epitope level resolution. On six diverse sets of antigen specific antibodies we demonstrate that SPACE2 accurately clusters antibodies that engage common epitopes and achieves far higher data set coverage than clonal clustering and SPACE1. Furthermore, we show that the functionally consistent structural clusters identified by SPACE2 are even more diverse in sequence, genetic lineage, and species origin than those found by SPACE1. These results reiterate that structural data improves our ability to identify antibodies that bind the same epitope, adding information to sequence-based methods, especially in data sets of antibodies from diverse sources. SPACE2 is openly available on Github (https://github.com/oxpig/SPACE2).

## Introduction

Antibodies are important components of the adaptive immune system. An antibody recognises foreign particles by binding to a specific site, the epitope, on their surface. As antibody function is tightly linked to the epitope it engages, studying epitopes is essential to understand immunology. For example, determining epitope specificities of antibody repertoires can increase our understanding of the immune response to disease (1, 2) or differences of the immune system between individuals (3). Furthermore, epitope profiling can be applied in antibody drug discovery to identify both new binders to a desired target (1, 4, 5) and binders with improved affinity (6). Epitopes can be determined at high resolution by solving the structure of an antibody in complex with its antigen. However, structure determination methods are too resource intensive to be used to explore large data sets (7). Experimental epitope binning methods, such as competition assays (8), scale better, but it remains difficult to analyse very large data sets as costs grow at O(n^2^) with the number of antibodies (n) to be evaluated. Competition assays also only offer low resolution as they struggle to distinguish between antibodies that bind distinct sites but that overlap sterically.

Prior computational clustering of antibodies into functional groups that engage the same epitope can reduce the number of experiments that need to be run or even remove the need for experimental epitope determination entirely. Most computational epitope profiling methods group antibodies based on sequence similarity. Clonotyping, the most widely used method, attempts to link antibodies that originate from the same progenitor B-cell (9, 10). The exact definition of a clonotype varies across the literature. Commonly antibodies that originate from the same heavy chain V and J genes, match in CDRH3 length and exceed a threshold CDRH3 sequence identity are considered a clonotype. Threshold values between 80 and 100% have been reported. To introduce additional leniency the requirement for matching J genes can be neglected (10). Clonal clustering is usually highly accurate, antibodies within a cluster tend to engage the same epitope.

Clonotyping was originally intended to trace lineages of antibodies within an individual. Its use in functional clustering thus makes the assumption that antibodies against a given epitope must originate from progenitor B-cells with shared genetic origins. However, antibodies from different lineages and with highly dissimilar sequences can adopt a similar binding site geometry and engage the same epitope (11–15). The ability to determine functional convergence is especially important when comparing the immune response of individuals, as different individuals exhibit personalised immunoglobulin gene usages (3). As clonotyping is not able to link antibodies from distinct genetic lineages, it loses power when analysing antibodies originating from different sources.

Alternative methods have been developed to try and identify functionally equivalent antibodies that are not similar in sequence. Clustering antibodies by sequence similarity across predicted paratope residues can link antibodies from different clonotypes (16). However, methods clustering antibodies based on structural similarity are even better suited to detect less related sequences with functional convergence (11, 12), because binding site structure provides more direct evidence of antibody function than its sequence.

In a previous study we described the SPACE1 method (11), which clusters antibodies based on structural similarity of homology models. The algorithm accurately clusters antibodies that bind the same epitope and is able to functionally link antibodies with diverse sequences. However, SPACE1 is limited by the coverage of homology modelling (in the original study only 73% of the data could be modelled to a usable standard) and its inaccuracies. The method was also only benchmarked at the level of domain consistency on one virus class. Recent progress in machine learning-based antibody structure prediction has led to more accurate structural models than those obtained with homology based approaches, especially in cases where no template with high sequence similarity is available (17–24). Higher accuracy and higher confidence in structural models also allow increased coverage and have the potential to improve structure-based epitope profiling.

Here, we present the Structural Profiling of Antibodies to Cluster by Epitope 2 (SPACE2) algorithm. SPACE2 builds on recent progress in machine learning-based antibody structure prediction and uses a novel clustering protocol systematically optimised and extensively benchmarked on epitope-resolution binding data. We show that SPACE2 outperforms SPACE1 by improving data coverage and identifying clusters even more diverse in sequences, genetic lineages and species origin. These results underline that structural data, which which can now be rapidly and easily generated through structure prediction tools, contains orthogonal functional information to sequence and should be considered when investigating antibody function.

## Methods

### Data sets

Six data sets of antigen specific antibodies were used to analyse SPACE2 clustering performance.

The training set on which the clustering algorithm, thresholds and antibody region were set consisted of 3051 antibodies against the SARS-CoV-2 receptor binding domain (RBD). Antibodies were annotated with groups of overlapping epitopes originating from mutation escape profiling (25). We refer to this dataset as the Cao et al. (25) training set throughout the paper.

CoV-AbDab (26), a data set of anti-lysozyme antibodies, a non-public data set of antibodies against Ebola viruses (EV) and two non-public data set of antibodies against non-viral targets (NVA1 and NVA2) were used as additional data sets to evaluate SPACE2. CoV-AbDab is a database of antibodies against coronavirus antigens, such as those from SARS-CoV-2, SARS-CoV-1 and MERS-CoV. A version of CoV-AbDab timestamped the 3^rd^ October 2022 was used containing 10,719 antibodies with sequence data. As CoV-AbDab is a collection of antibodies reported in the literature it contains the Cao et al. (25) training set. When using CoV-AbDab as a test set (denoted as CoV-AbDab (test)) the training set was removed and only the remaining 7685 antibodies included. Epitope data in CoV-AbDab is reported as in the original publications and ranges from antigen to domain level.

A data set of anti-lysozyme antibodies was created from all 53 lysozyme specific antibodies in the structural antibody database (SAbDab) (27, 28) for which the antibody-antigen complex structure has been solved. Antibodies were grouped by their epitope using the Ab-ligity method (12) and annotated as binding to the same epitope if their Ab-ligity score was greater than a threshold of 0.1 (as in the original paper). Similarity of epitopes within an epitope group was confirmed by visual inspection.

The EV set contains 126 antibodies with epitope data ranging from antigen to domain level. The NVA1 set contains 31 antibodies with epitope data from competition assays. NVA2 contains 33 antibodies with epitope data from mutation escape profiling.

### SPACE1

The original SPACE1 method clusters antibodies by the structural similarity of homology models. The algorithm was run as detailed in Robinson et al. (11).

Homology models were produced using ABodyBuilder (29). ABodyBuilder uses structures from a database to build its models. In this study we used quality-filtered SAbDab (27, 28) entries timestamped before 6^th^ July 2022. Quality filtering restricts structures to those solved by X-ray crystallography, and excluded structures with a resolution of >2.5 Å and structures containing residues with a B-factor >80. In a standard ABodyBuilder run the method first attempts to model CDR loops with a template database search method (30). If no suitable template is found for a CDRs, hybrid homology/*ab initio* modelling is performed (29). Only models for which homology templates for all six CDR loops were found are used for clustering in the SPACE1 method to keep the models as accurate as possible.

The remaining homology models are clustered by structural similarity of CDRs. Models are split into groups of antibodies with identical CDR lengths. Antibodies in each group are then clustered using a greedy clustering algorithm. The first antibody in the group was selected as the cluster centre and all antibodies with a CDR C_*α*_ RMSD smaller than a specified threshold after alignment of framework residues were added to the cluster. After all antibodies have been compared against the first cluster centre the algorithm selects the next unclustered antibody as a new cluster centre and cluster members are chosen as in the previous step. In addition to the RMSD threshold of 0.75 Å suggested by Robinson et al. (11), we also assessed the performance at a 1.25 Å threshold.

### SPACE2

Our novel SPACE2 algorithm clusters antibodies by the similarity of models obtained from an ML-based structure prediction tool. The method functions in four main steps. Initially, a structural model of the antibody Fv is produced using ABodyBuilder2 (20). ABodyBuilder2 is a deep-learning based tool for antibody structure prediction and was trained on SAbDAb structures timestamped up to 31^st^ July 2021. The models are then split into groups of identical CDR lengths. Models in each group are then structurally aligned on the C*α* of residues in framework regions and a pairwise distance matrix is computed of the C*α* RMSDs of CDR loop residues. The antibodies are then clustered based on these distances.

#### Clustering algorithms

Eight different clustering algorithms were explored (agglomerative clustering, affinity propagation, DBSCAN, OPTICS-xi, OPTICS-DBSCAN, K-means, Butina clustering, greedy clustering). Agglomerative clustering (31), affinity propagation (32), DBSCAN (33), OPTICS-xi, OPTICS-DBSCAN (34) K-means (35) were performed using the scikit-learn (36) implementation. Butina clustering (37) was preformed using the RDKit implementation (38). A greedy clustering algorithm, grouping antibodies as the algorithm described in section, was implemented.

Parameters and evaluated ranges for each algorithm are shown in Table S1. The K-means algorithm requires an additional parameter (K) which corresponds to the predetermined number of clusters. K was set to the number of clusters obtained from agglomerative clustering using the best-performing parameters.

#### SPACE2-HC

A variation of the SPACE2 algorithm was implemented that clusters antibodies based on the structural similarity of heavy chains only (SPACE2-HC). The light chains were included for the modelling step, as ABody-Builder2 (20) requires sequences of both chains as an input. After this step, light chains were ignored. Antibodies were grouped based on the length of the heavy chain CDRs, aligned on heavy chain framework regions, and the C_*α*_ RMSD of CDRs H1-3 calculated. Agglomerative clustering with a ‘complete’ linkage criterion was used as the clustering algorithm of SPACE2-HC.

#### SPACE2-Paratope

A second variation of the SPACE2 algorithm was implemented that clusters antibodies based on the structural similarity of CDR loops which are predicted to form part of the paratope (SPACE2-Paratope). Structural models were produced with ABodyBuilder2 (20). The Paragraph method (39) with a classifier cut-off of 0.734, as suggested in the original paper, was then used to predict residues that are part of the paratope based on the models. All CDRs containing at least one paratope residues were then labelled as paratope CDRs. Antibodies were divided into groups containing the same set of paratope CDRs. Antibodies in each group were further grouped based on the length of the paratope CDRs, aligned on heavy chain framework regions, and clustered based on the C_*α*_ RMSD of paratope CDRs. Agglomerative clustering with a ‘complete’ linkage criterion was used as the clustering algorithm of SPACE2-Paratope.

### Numbering scheme and region definitions

IMGT numbering (40) and North CDR definitions (41) are used throughout.

### Analysis of structural clusters

#### Domain/Epitope consistent clusters

Antibody clusters generated for the Cao et al. (25) training set, the NVA1 set, the NVA2 set and the anti-lysozyme set were classified as ‘epitope-consistent’ or ‘epitope-inconsistent’. ‘Epitope-consistent’ clusters of the Cao et al. (25) training, NVA1 and NVA2 sets only contain antibodies that bind to the same epitope group as determined by experimental epitope binning. ‘Epitope-consistent’ clusters of the lysozyme data set only contain antibodies that bind to the same residue-level epitope determined using crystal structures.

Owing to the lower resolution of epitopes reported in the EV set and CoV-AbDab, clusters of these data sets were classified as ‘domain-consistent’ and ‘domain-inconsistent’. EV set clusters were labelled as ‘domain-consistent’ if they only contain antibodies that engage the same antigen domain. CoV-AbDab clusters that satisfy the following rules, consistent with previous studies (11), were determined to be ‘domain-consistent’:

1. clusters that only contain antibodies that bind to the same antigen and domain
2. clusters that contain antibodies binding to the same domain and others that bind to the same antigen without domain-level resolution
3. clusters that only contain antibodies that bind to the same antigen but do not have domain-level resolution of epitope data
4. clusters with internally consistent epitope data, e.g. a cluster of antibodies labelled to bind to the spike (S) protein N-Terminal Domain (NTD) and others labelled as S non-RBD binders, as S NTD is a subdomain of S non-RBD.

#### Performance metrics

Through out this study we use seven metrics to analyse functional clustering. Two accuracy metrics, the fraction of epitope-consistent clusters (number of epitope-consistent multiple-occupancy clusters / number of multiple-occupancy clusters) and the fraction of clustered antibodies in epitope-consistent clusters (number of antibodies in epitope-consistent multiple-occupancy clusters / number of antibodies in multiple-occupancy clusters) were used. Two coverage metrics, the number of multiple-occupancy clusters and the number of antibodies in multiple-occupancy clusters were used. In order to examine accuracy and coverage with one measure we also calculated the number of antibodies in consistent multiple-occupancy clusters. Two further metrics were used to assess the diversity of antibodies within clusters, the fraction of functionally consistent clusters containing antibodies from more than one clonotype as well as the mean CDRH3 sequence identity within functionally consistent clusters.

#### Random baseline

Random clustering was performed as a baseline. The distribution of cluster sizes obtained from the evaluated clustering algorithm with specific parameters was recorded. Clusters with an identical size distribution were then sampled randomly from the data set and performance metrics were calculated. Sampling was repeated 100 times and the metrics averaged.

#### Clonotyping

Clonotyping was performed using an in-house script. Lenient VH-clonotyping and Fv-clonotyping threshold conditions by community standards were used (9, 10). A VH-clonotype was defined as matching IGHV gene, length matched CDRH3 and >80% CDRH3 sequence similarity. Fv-clonotypes were defined as matching VH-clonotype, matching of IG[K/L]V genes, length matched CDRL3 and >80% sequence identity of CDRL3.

## Results

The original SPACE1 algorithm was developed to cluster antibodies by structural similarity with the aim of better identifying functional convergence. It grouped antibodies based on the structural similarity of homology models. This method was not systematically optimised and only benchmarked on a single data set of low resolution epitope data. Newly available ML-based structure prediction tools produce more accurate models and have better coverage than homology modelling. Here, we introduce SPACE2 which uses a state-of-the-art antibody structure prediction method and a novel clustering protocol that has been extensively optimised and then benchmarked on several data sets of high resolution epitope data.

SPACE2 clusters antibodies in four main steps. Initially, structural models are produced using ABodyBuilder2 (20). Models are then separated into groups of antibodies with identical lengths of the six CDRs, followed by the computation of a pairwise distance matrix of CDR C_*α*_ RMSDs. In the final step, a clustering algorithm divides the antibodies into structural clusters. Optimisation of the clustering protocol was performed on a training set of 3051 antibodies against the SARS-CoV-2 receptor binding domain (RBD) (25).

### Evaluating an optimal clustering algorithm

We tested eight widely used clustering algorithms, greedy clustering, affinity propagation (32), Butina clustering (37), DBSCAN (33), OPTICS-DBSCAN, OPTICS-xi (34), agglomerative clustering (31) and K-means (35), for their ability to correctly group functionally consistent antibodies in the Cao et al. (25) training set. To assess the methods we used the number of antibodies in epitope-consistent multiple-occupancy clusters as our target performance metric as it provides a trade-off between clustering accuracy and data set coverage. High accuracy or coverage metrics individually do not necessarily indicate a good epitope profiling method. High accuracy can be achieved by dividing the data set into very small clusters that are highly likely to be epitope/domain-consistent but do not cover the full diversity of antibodies able to engage a given epitope. Maximal coverage can be achieved by putting all antibodies into a single cluster, which does not provide any useful epitope information.

A parameter scan was carried out to find the optimal setting for each clustering method. Evaluated parameter ranges and optimal values are shown in Table S1. As expected lenient parameters increased data set coverage whereas stringent parameters improve accuracy and the trade-off was maximised at intermediate values. The best performing algorithms, as defined by maximising the number of antibodies in epitope-consistent multiple-occupancy clusters, were agglomerative clustering (optimal parameters: linkage criterion = complete, RMSD distance threshold = 1.25 Å), OPTICS-xi (optimal parameters: xi ≤ 0.01, RMSD distance threshold = 2 Å) and K-Means (optimal parameters: initialisation method = K-means++) where K was set to the number of clusters obtained by agglomerative clustering with optimal parameters (Fig. 1). As K-means does not lead to an improvement over agglomerative clustering, it was disregarded for further analysis.

**Fig. 1.**
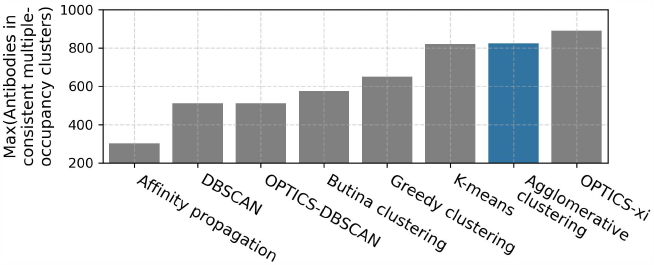
Examination of clustering algorithms. Parameter scans of eight clustering algorithms were performed using the Cao et al. (25) training set. The performance of the clustering was measured in terms of the number of antibodies in epitope-consistent multiple-occupancy clusters (y-axis). The maximum value of this metric achieved by a specific algorithm across all evaluated parameters when clustering the Cao et al. (25) training set is shown. Evaluated parameter ranges and optimal values are shown in Table S1. The agglomerative clustering algorithm selected for SPACE2 is highlighted in blue.

Agglomerative and OPTICS-xi clustering were compared in more detail (Table S2). Both algorithms achieve a similar clustering accuracy and data set coverage. Agglomerative clustering produces larger clusters with a mean cluster size of 3.0 members and a maximum of 28 than OPTICS-xi clusters with mean 2.7 and maximum 11. When epitope consistent clusters are larger it suggests they are better capturing the full diversity of the antibodies able to engage a given epitope. Therefore, agglomerative clustering was selected for use in SPACE2.

### Examining the behaviour of agglomerative clustering across different structural similarity thresholds

The RMSD threshold parameter of agglomerative clustering determines the leniency of the algorithm as it sets the maximum distance between any two antibodies in a cluster. Small thresholds restrict clustering to highly similar structures, whereas larger values allow clusters to contain more dissimilar antibodies. We evaluated agglomerative clustering for threshold values between 0.5 and 5 Å to assess how clustering results are affected.

Four metrics were monitored to assess the accuracy of clustering and the data set coverage. The fraction of epitope-consistent clusters (number of epitope-consistent multiple-occupancy clusters / number of multiple-occupancy clusters) and the fraction of clustered antibodies in epitope-consistent clusters (number of antibodies in epitope-consistent multiple-occupancy clusters / number of antibodies in multiple-occupancy clusters) were used as an accuracy measure. The number of multiple-occupancy clusters and the number of antibodies in multiple-occupancy clusters provide information on data set coverage.

Clustering accuracy and data coverage show a strong dependence on the RMSD threshold (Fig. 2). At thresholds ≤ 0.75 Å the clustering is highly accurate. More than 80% of clusters are epitope-consistent and approximately 80% of clustered antibodies are in epitope-consistent clusters. Increasing the threshold leads to a rapid drop in accuracy but improves data set coverage. The number of antibodies in multiple occupancy clusters starts to plateau at around 3 Å. The large changes of accuracy and data coverage as a function of threshold suggest that the threshold should be adjusted depending on the aim of the epitope profiling task. Optimal clustering is achieved at a value of 1.25 Å, as defined by maximising the number of antibodies in epitope-consistent multiple-occupancy clusters. However, the threshold can be set to any value between 0.75 and 3 Å to increase accuracy or coverage.

**Fig. 2.**
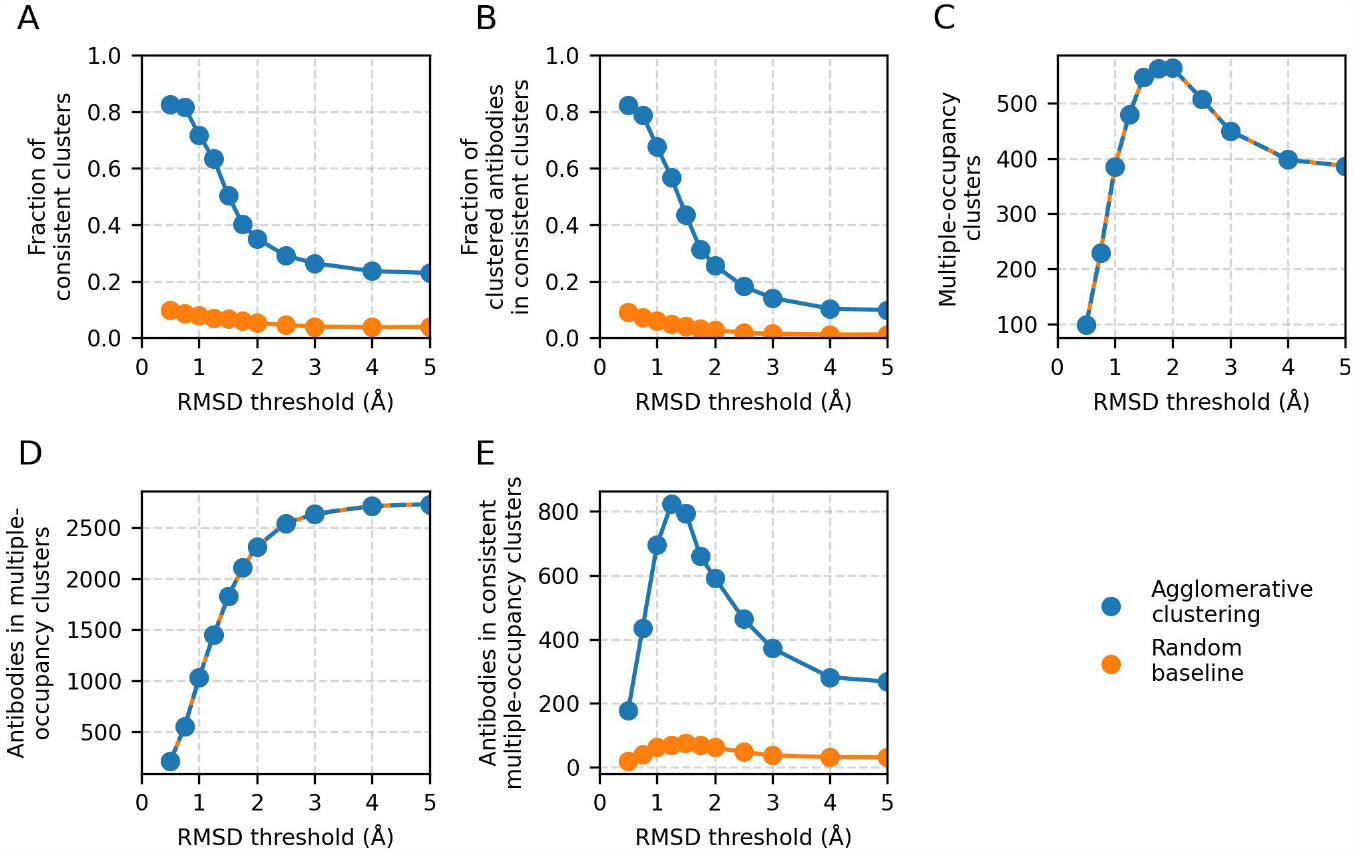
Results of agglomerative clustering as a function of RMSD threshold on the Cao et al. (25) training set. Agglomerative clustering with a ‘complete’ linkage criterion was performed for threshold values between 0.5 Å and 5 Å. The values of the five performance metrics are plotted against evaluated threshold values: (A) fraction of epitope-consistent clusters, (B) fraction of clustered antibodies in epitope-consistent clusters, (C) number of multiple-occupancy clusters, (D) number of antibodies in multiple-occupancy clusters, (E) number of antibodies in epitope-consistent multiple-occupancy clusters. Results of a random clustering baseline (see methods) are shown for comparison. Values for the number of multiple-occupancy clusters and antibodies in multiple-occupancy clusters for the random baseline are matched to agglomerative clustering.

In all the analysis to this point we have reported only on clusters that are 100% epitope-consistent (i.e. only contain antibodies against the same epitope). To measure the inconsistency of the remaining clusters, we analysed the fraction of clusters in which at least 70% of the antibodies engage the same epitope. An additional 12% of clusters are >70% epitope-consistent and these clusters contain an extra 26% of all antibodies contained in multiple-occupancy clusters (Fig. S1). This result indicates that even those clusters our standard performance metrics are marking as incorrect may contain large amounts of useful information.

### Evaluating the optimal region for clustering

The SPACE2 method calculates structural similarity of antibodies across all six CDRs. However, not all CDRs are equally involved in binding and we expect the structure of some CDRs to be more important in determining epitope specificity than the structure of others. Therefore, we investigated how the choice of CDRs over which RMSDs are calculated impacts clustering. We assessed two variations of SPACE2 that cluster based on subsets of CDRs.

In the first variation, the algorithm was adapted to consider structural similarity of heavy chain CDRs only (SPACE2HC). This approach was motivated by sequence-based methods such as clonotyping which often achieve good performance considering only the heavy chain sequence. In SPACE2-HC antibodies were grouped based on the length of the heavy chain CDRs, aligned on heavy chain framework regions, C_*α*_ RMSD of CDRs H1-3 calculated and clustered with an agglomerative clustering algorithm with a ‘complete’ linkage criterion. An RMSD threshold of 1.25 Å was found to optimise SPACE2-HC (Fig. S2). SPACE2-HC performed worse than the standard SPACE2 algorithm as measured by a 33% drop in the trade-off metric of antibodies in epitope-consistent clusters (Table S3). While SPACE2-HC slightly increased data set coverage, a substantial decrease in accuracy was observed.

A second variation of SPACE2 was implemented to cluster antibodies based only on the similarity of CDR loops that contain paratope residues (SPACE2-Paratope). Paratope residues were predicted using the Paragraph method (39). Models were grouped by the combination of CDR loops that contain paratope residues (paratope CDRs). Models were then grouped again based on the length of paratope CDRs, aligned on framework regions and the C_*α*_ RMSD of paratope CDRs was calculated. A RMSD threshold of 1.5 Å was found to optimise agglomerative clustering for SPACE2-Paratope (Fig. S2). Measured by the trade-off metric, SPACE2-Paratope performed worse than standard SPACE2 (Table S3). A slight drop in both clustering accuracy and data coverage was observed.

The best clustering results were achieved by clustering based on the structural similarity of all six CDR loops. Therefore, the standard SPACE2 methods was chosen as the clustering protocol for further analysis.

### SPACE2 performs well on sets of antibodies against diverse targets

SPACE2 was tested on five data sets of antigen specific antibodies using the clustering algorithm (agglomerative clustering) and parameter choices (complete linkage criterion, 1.25 Å RMSD threshold) defined on the Cao et al. (25) training set. The test sets comprised a data set of anti-lysozyme antibodies, a non-public data set of anti-Ebola virus antibodies (EV), two non-public data sets of antibodies against non-viral targets (NVA1 & NVA2) and CoV-AbDab (test), a version of CoV-AbDab with training set overlap removed (see methods) (26). An overview of results from clustering the test sets is shown in Table 1.

**Table 1.**
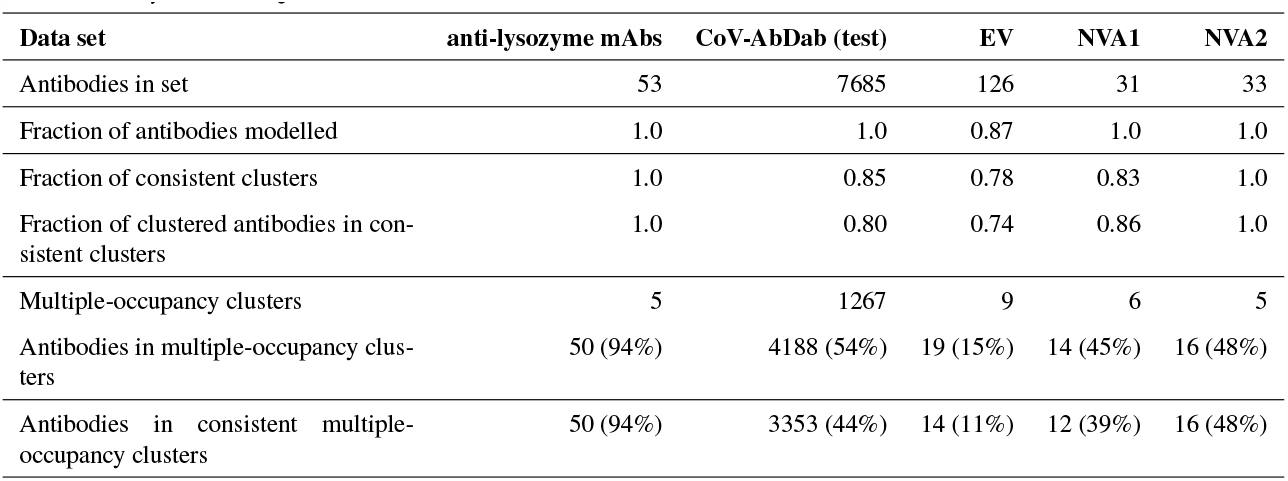
Performance of SPACE2 on test data sets. Values of the five performance metrics as well as the fraction of antibodies successfully modelled with ABodyBuilder2 are shown for each data set. For the two metrics of number of antibodies in multiple-occupancy clusters and number antibodies in epitope-consistent multiple-occupancy clusters additionally a percentage is shown, indicating the percentage of antibodies in the data set. CoV-AbDab (test) denotes the subset of CoV-AbDab that is not contained in the Cao et al. (25) training set (see methods). CoV-AbDab (test) was used for this analysis to prevent testing on training set antibodies.

The anti-lysozyme data set contains antibodies against five distinct epitopes. SPACE2 clusters antibodies in this set with high accuracy as 100% of clusters are epitope-consistent. Good data coverage is observed and 50 of the 53 antibodies fall into multiple-occupancy clusters. SPACE2 divides the data set into eight clusters (Fig. 3). We observe three cases where antibodies binding a common epitope are separated into two clusters. Looking at these cases in more detail shows that despite engaging the same epitope the antibody structures do not overlay perfectly. In each case, we observe antibodies that bind the epitope in two different binding poses and these are separated into distinct clusters by SPACE2. These results show that SPACE2 groups antibodies with a high resolution.

**Fig. 3.**
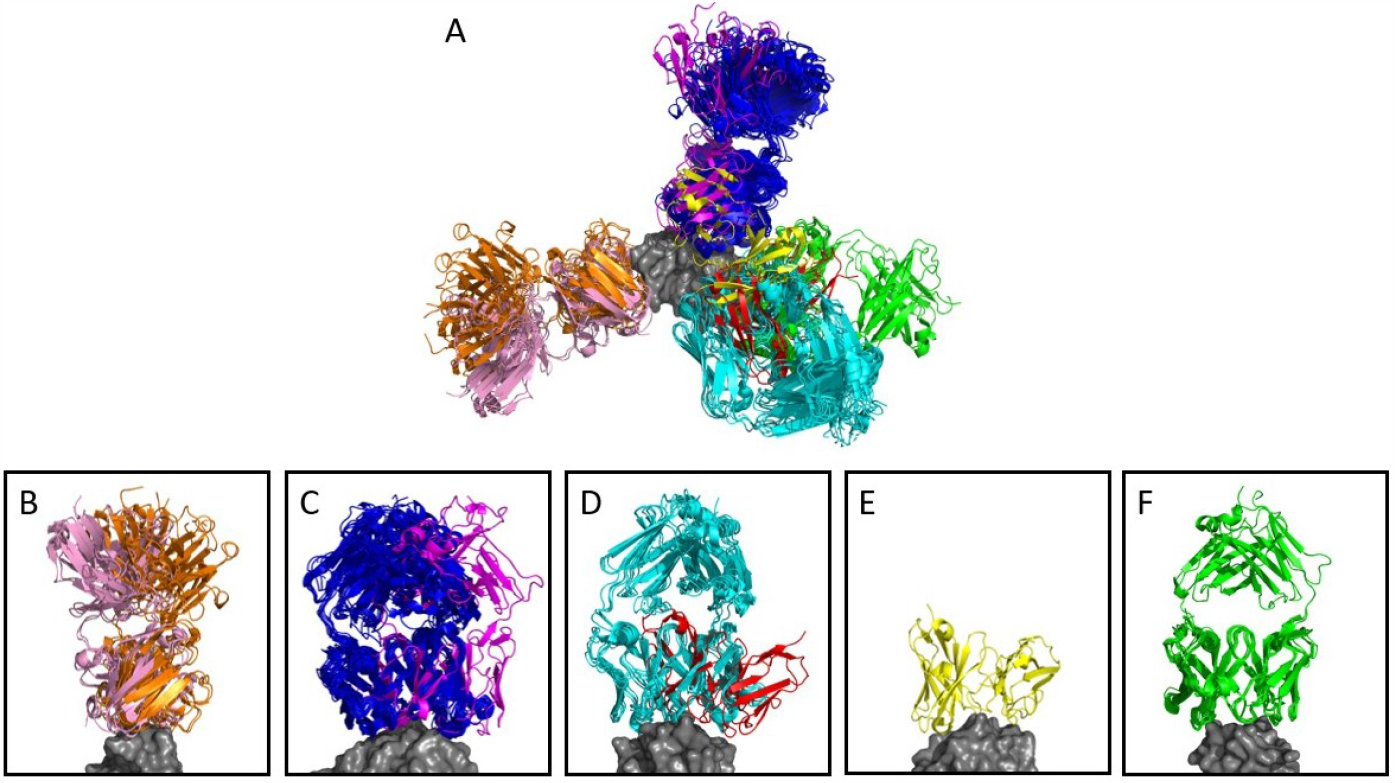
Anti-lysozyme antibodies. Crystal structures of 53 antibody-lysozyme complexes are shown aligned on the antigen structure (grey). Antibodies are coloured according to the clusters assigned by SPACE2. (A) Overlay showing all 53 antibody-lysozyme complexes. (B-F) Each panel shows all antibodies that bind to one of the five lysozyme epitopes as defined bu Ab-ligity (12). Panels B, C and D each contain two sets of antibodies that do not overlay perfectly indicating a difference in binding pose. SPACE2 separates antibodies binding the same epitope in a different binding pose into distinct clusters as indicated by the colouring.

SPACE2 also achieves a high clustering accuracy on CoV-AbDab (test), the EV set, the NVA1 set and the NVA2 set. 85%, 78%, 83% and 100% of clusters in the four sets are domain/epitope-consistent, respectively. Domain/epitope-consistent clusters comprise 80%, 74%, 86% and 100% of all antibodies grouped into multiple-occupancy clusters.

Data coverage differs for the four data sets (see methods for definition of coverage metrics). Coverage of CoV-AbDab (test), NVA1 and NVA2 is high with 54%, 45% and 48% of all antibodies contained within multiple-occupancy clusters. In comparison, only 19 of 126 EV set antibodies are grouped into multiple-occupancy clusters. The EV set is relatively small and contains antibodies against the Ebola virus glycoprotein, a large multi-domain protein with many potential epitopes (42). We do not expect to observe many antibodies engaging the same residue-level epitope in a small data set of antibodies against a target with many epitopes, which is likely why we see low coverage.

Overall, SPACE2 generalises well to the test sets. The algorithm achieves a high clustering accuracy on all five data sets and a good coverage on the CoV-AbDab (test), NVA1, NVA2 and the anti-lysozyme database. Coverage of the EV set is comparably low, indicating a challenge in clustering smaller data sets of epitope diverse antibodies.

### Advances in structure prediction improve structure based computational epitope profiling

We compared the performance of SPACE2 to SPACE1, our previous structural epitope profiling method. SPACE1 (11) groups antibodies based on structural similarity of homology models produced with ABodyBuilder (29) followed by greedy structural clustering at an RMSD threshold of 0.75 Å.

We once again used the number of antibodies in epitope/domain-consistent multiple-occupancy clusters which balances both accuracy and coverage as our metric for comparing performance. SPACE2 outperforms SPACE1 using its suggested threshold (RMSD threshold 0.75 Å) (Table S4). As SPACE2 uses an RMSD threshold of 1.25 Å, we also explored a range of RMSD values to see whether the difference in RMSD threshold is the driver for the difference in performance. We found that a threshold of 1.25 Å improved SPACE1 clustering (Table S4) but it was still significantly worse than SPACE2. SPACE1 with a 1.25 Å threshold results in an 18% and 9% decrease in antibodies in epitope/domain-consistent multiple-occupancy clusters on the two largest data sets, CoV-AbDab and the Cao et al. (25) training set, respectively (Table 2). SPACE2’s better performance is driven by better coverage while achieving a similar accuracy. This increase in coverage arises due to improvements in the modelling step, ML-based structure prediction allows nearly all of the antibodies to be modelled (Table 2), higher model quality as well as the optimised clustering protocol.

**Table 2.** Comparison of SPACE2, SPACE1 and clonotyping. The original SPACE1 algorithm was evaluated at an RMSD threshold of 1.25 Å. Two protocols were used for clonotyping (see methods). VH-clonotyping is restricted to genes and sequence of the heavy chain. Fv-clonotyping considers both heavy and light chains. For the two metrics of number of antibodies in multiple-occupancy clusters and number antibodies in epitope-consistent multiple occupancy clusters additionally a percentage is shown, indicating the percentage of antibodies in the data set. The most important performance metric to consider when comparing different epitope profiling methods is the number of antibodies in epitope/domain-consistent multiple-occupancy clusters as high accuracy or coverage metrics individually may not indicate good performance. The fraction of epitope/domain-consistent clusters containing more than one VH-clonotype and the mean CDRH3 sequence identity observed within epitope/domain-consistent clusters are also given. The best result for each metric is highlighted in bold.

| Data set | CoV-AbDab |  |  |  | Training set |  |  |  |
| --- | --- | --- | --- | --- | --- | --- | --- | --- |
| Method | SPACE2 | SPACE1 | Clonotyping |  | SPACE2 | SPACE1 | Clonotyping |  |
|  |  |  | VH | Fv |  |  | VH | Fv |
| Fraction of antibodies modelled | **1.0** | 0.71 | - | - | **1.0** | 0.7 | - | - |
| Fraction of consistent clusters | 0.87 | 0.87 | 0.98 | **0.99** | 0.63 | 0.64 | 0.84 | **0.83** |
| Fraction of clustered antibodies in consistent clusters | 0.82 | 0.81 | 0.97 | **0.99** | 0.57 | 0.58 | 0.75 | **0.79** |
| Multiple-occupancy clusters | **1811** | 1165 | 1191 | 970 | **480** | 314 | 361 | 303 |
| Antibodies in multiple-occupancy clusters | **6271**<br>(59%) | 4010<br>(37%) | 4045<br>(38%) | 2754<br>(26%) | **1446**<br>(47%) | 935<br>(31%) | 1060<br>(35%) | 793<br>(26%) |
| Antibodies in consistent multiple-occupancy clusters | **5126**<br>(48%) | 3255<br>(30%) | 3916<br>(37%) | 2733<br>(25%) | **823**<br>(27%) | 538<br>(18%) | 797<br>(26%) | 628<br>(21%) |
| Fraction of clusters containing >1 VH-clonotypes | **0.81** | 0.71 | 0 | 0 | 0.55 | **0.58** | 0 | 0 |
| Mean CDRH3 sequence identity | **0.54** | 0.57 | 0.88 | 0.88 | **0.66** | 0.67 | 0.86 | 0.87 |

### SPACE2 improves coverage compared to clonotyping

Clonotyping is the most commonly used epitope profiling method. It clusters antibodies based on sequence similarity. As clonotyping assumes that antibodies against a given epitope must originate from progenitor B-cells with shared genetic origins it cannot detect functional convergence. Thus, the method is limited when clustering data sets of antibodies from different individuals or species. Here, we compare SPACE2 to two lenient clonotyping protocols, VH and Fv-clonotyping (see methods), on the two largest data sets which contain antibodies from diverse sources. The Cao et al. (25) training set consists of antibodies isolated from 165 human patients (25) and CoV-AbDab contains antibodies from a range of studies (∼450) and several species (26).

The performance of SPACE2 and the two clonotyping protocols are shown in Table 2. SPACE2 outperforms both VH and Fv-clonotyping in the key metric of antibodies in epitope/domain-consistent clusters on both data sets. Improvement in this metric is driven by increased data set coverage by SPACE2. We observe 33% and 21% more antibodies in multiple-occupancy clusters for Fv-clonotyping of CoV-AbDab and the Cao et al. (25) training set, respectively. Data coverage by VH-clonotyping is better but still substantially lower than SPACE2. On the other hand, SPACE2 is less accurate than both clonotyping protocols. However, the increase in coverage achieved by SPACE2 exceeds the drop in accuracy.

### SPACE2 identifies functional convergence signals

We next analysed the diversity of antibodies within the SPACE2 clusters of the Cao et al. (25) training set and CoV-AbDab, to see if we were identifying functionally similar antibodies with very different sequences.

The majority of SPACE2 clusters contain antibodies belonging to several clonotypes, highlighting the ability to link antibodies from different genetic lineages. Specifically, 55% of epitope-consistent clusters from the Cao et al. (25) training set and 81% of domain-consistent clusters from CoV-AbDab contain antibodies from more than one VH-clonotype (Table 2).

Moreover, we investigated the sequence similarity of antibodies within epitope/domain-consistent clusters. Clonotyping is limited to linking sequence similar antibodies as the method uses a CDRH3 sequence identity cut-off to cluster antibodies. Here, we use a lenient cut-off of 80%. We observe a mean CDRH3 sequence identity of 86% within epitope-consistent VH-clonotypes of the Cao et al. (25) training set and 88% for domain-consistent VH-clonotypes of CoV-AbDab. In comparison, SPACE2 clusters tend to be highly diverse in sequence. Epitope/domain-consistent clusters have a mean CDRH3 sequence identity of 54% and 66% for CoV-AbDab and the Cao et al. (25) training set, respectively (Table 2). A large number of CoV-AbDab clusters were observed with mean sequence identity below 40% (Fig. S3) and some clusters even contain pairs of antibodies with no common CDRH3 residues.

Structural clustering is also able to functionally link antibodies from different organisms. For CoV-AbDab, SPACE2 produced 26 functionally consistent clusters containing antibodies from more than one species and was able to group antibodies from human, mouse and rhesus macaque origins. In comparison, optimised SPACE1 was only able to detect 18 domain-consistent inter-species clusters and clonotyping is unable to link antibodies from different species.

#### SPACE2 informs on functional convergence of sequence dissimilar antibodies

We examined in more detail a SPACE2 cluster of the CoV-AbDab data set with 12 members (368.07.C.0221, BD55-4342, BD55-5339, BD55-5550, BD55-5856, BD55-6024, BD55-6223, BD55-6372, BD55-6596, BD57-074, C018, EY6A) (Fig. 4). Eleven of the antibodies engage the spike protein RBD, the final member is annotated as a spike protein binder with unknown domain. Clustering by SPACE2 suggests that these antibodies, determined to bind to the same domain of the spike protein, engage the same residue-level epitope.

**Fig. 4.**
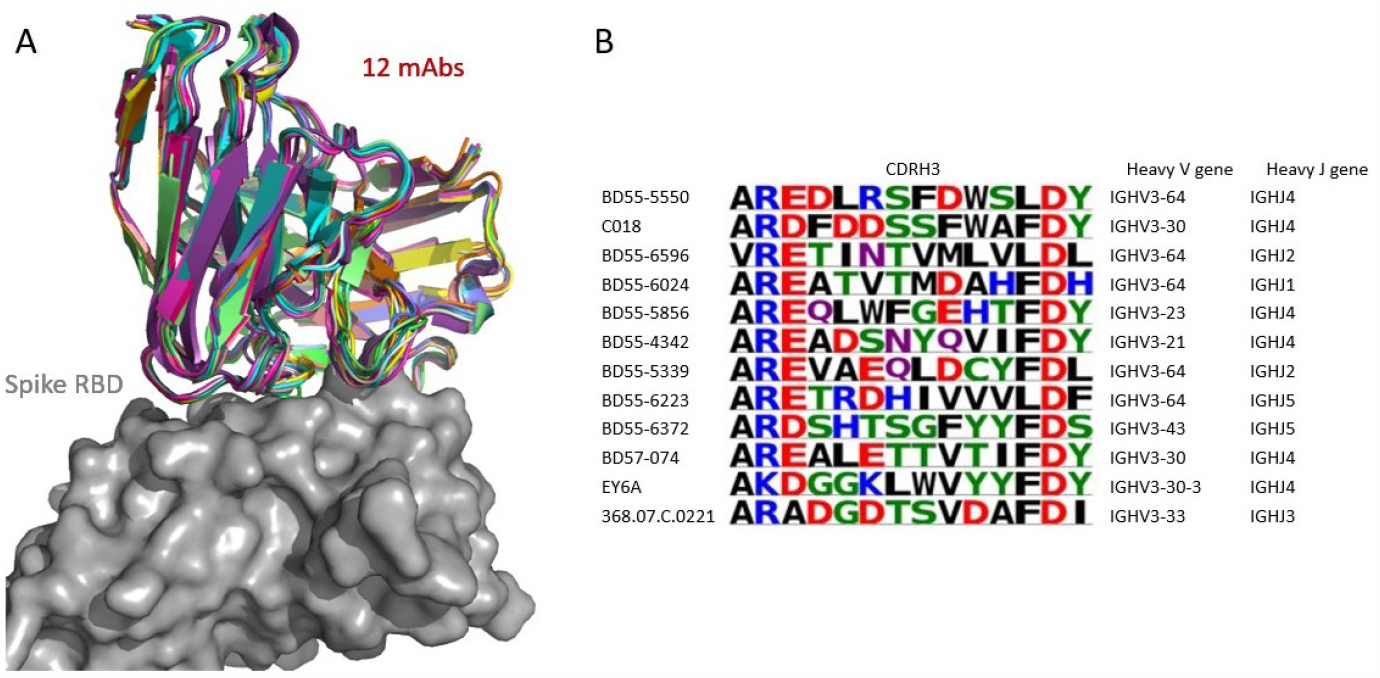
SPACE2 identifies functional convergence of antibodies. Twelve membered CoV-AbDab cluster (368.07.C.0221, BD55-4342, BD55-5339, BD55-5550, BD55-5856, BD55-6024, BD55-6223, BD55-6372, BD55-6596, BD57-074, C018, EY6A) with a mean CDRH3 sequence identity of 33%. The crystal structure of EY6A in complex with its antigen is available (PDB 6ZCZ), SPACE2 suggests that the 11 remaining antibodies bind to the same residue-level epitope. (A) Structural models of the 12 members are overlaid with the crystal structure of EY6A (not shown) in complex with the spike protein RBD (grey). (B) CDRH3 sequence alignment of all 12 cluster members coloured by chemical properties of amino acid residues (produced with Logomaker (43)) and heavy chain V and J genes.

The 12 antibodies are highly diverse in sequence and genetic lineage. The cluster shows a mean CDRH3 sequence identity of 33%. The antibodies possess a CDRH3 of length 12 and 8 of these residues differ on average. The most distant pair of antibodies in the cluster are BD55-6596 and EY6A which differ in 11 of 12 CDRH3 residues. The 12 antibodies originate from 7 different IgGH genes and fall into 12 separate VH-clonotypes.

The improvement of SPACE2 over SPACE1 can be seen when examining how these antibodies were clustered by SPACE1. Using SPACE1 with an optimised threshold only six of these antibodies (BD55-6024, BD55-6223, BD55-6596, BD57-074, C018, EY6A) were grouped together and even these six were part of a larger functionally inconsistent cluster with 44 members. BD55-5339 was in a separate functionally inconsistent SPACE1 cluster with 4 members and the remaining five antibodies were not placed in multiple-occupancy clusters.

#### SPACE2 identifies epitopes targeted by multiple species

As the CoV-AbDab database contains antibodies from multiple species (human, mouse, rhesus macaque) we examined whether SPACE2 can identify epitopes targeted by multiple species. There were 26 SPACE2 clusters of the CoV-AbDab database that contained antibodies from more than one species. We examined a SPACE2 cluster with seven members (368.02a.C.0049, B13, BD55-6574, BD57-092, DK15, Fab-160, SW186) (Fig. 5). The cluster contains six antibodies that engage the spike protein RBD and one spike specific antibody without domain-level epitope data. Five of the antibodies are human and originate from human patients, phage-display and transgenic mice. The remaining are murine and were found in immunised mice. SPACE2 suggests that these human and mouse antibodies engage the same residue-level epitope which highlights its ability to detect public epitope targeted by multiple species.

**Fig. 5.**
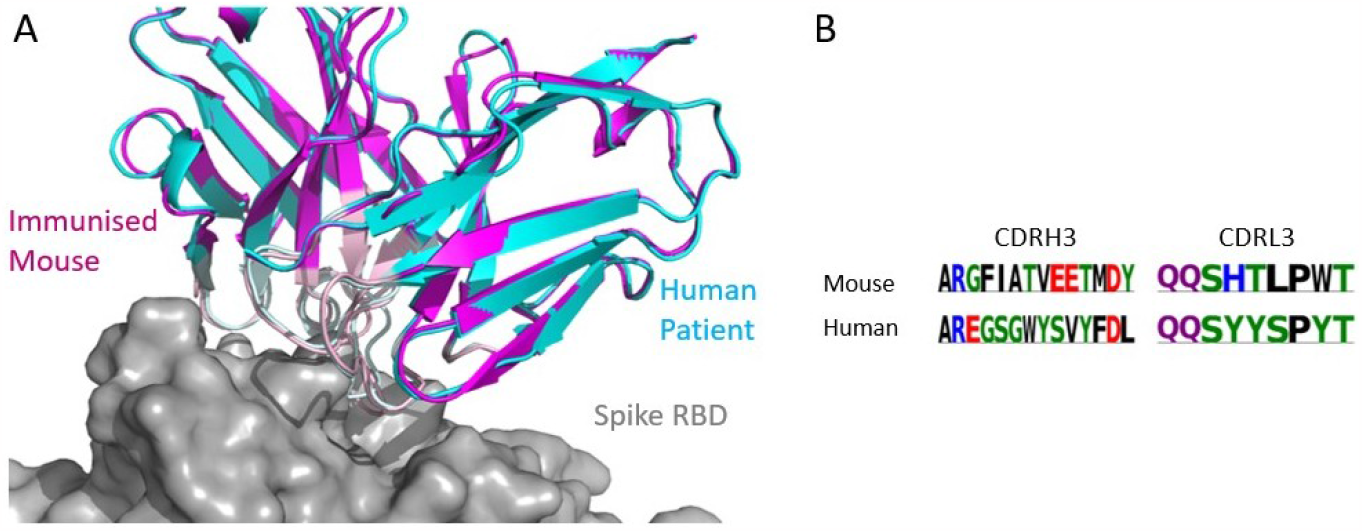
SPACE2 identifies epitopes targeted by multiple species. A SPACE2 cluster from CoV-AbDab containing antibodies from multiple species. Two representatives from a cluster comprising murine (SW186) and human (BD57-092) antibodies are shown. The crystal structure of SW186 is available (PDB 8DT3), SPACE2 suggests that BD57-092 binds to the same residue-level epitope. (A) The structural models of SW186 (pink) and BD57-092 (cyan) were overlaid with the crystal structure of SW186 (not shown) in complex with the spike protein RBD (grey). CDR regions of both antibodies are highlighted by the lighter colouring. (B) CDRH3 sequence alignment of the two antibodies coloured by chemical properties of amino acid residues (produced with Logomaker (43)).

SPACE1 with an optimised threshold was only able to link one of the two murine antibodies in this cluster to human structures, while clonotyping is unable to link mouse and human antibodies due to different gene usage.

## Discussion

Accurately identifying the epitope of antibodies is a key step in understanding immunology and in the design of new biologic drugs. Such data are conventionally determined experimentally either by solving individual antibody-antigen crystal structures or by epitope binning methods such as competition binding assays. Prior computational clustering of antibodies into functional groups could reduce the number of experiments that need to be carried out or even remove the need for them entirely. Clonal clustering is most commonly used for this purpose, where antibodies are grouped by sequence identity and genetic lineage. But these types of methods will miss antibodies with low sequence identity that have functionally converged and target common epitopes (11–15, 44). In a previous study, we reported the Structural Profiling of Antibodies to Cluster by Epitope (SPACE1) method (11), which clusters antibodies by structural similarity of their homology models. This method showed that structure-based epitope profiling is better able to detect the full breadth of functional convergence. However, SPACE1 was limited by the coverage of template-based modelling and its inaccuracies. The method was also only benchmarked at the level of domain-consistency of antibodies against one virus class. Here, we introduce SPACE2, an updated method which uses the latest machine learning-based antibody structure prediction technology (20) and a novel clustering protocol systematically optimised on epitope-resolution data.

We show across six data sets that SPACE2 can accurately bin antibodies that engage the same epitope and achieve high data coverage. Available crystal structures of antibody-antigen complexes reveal that SPACE2 tends to group antibodies that bind to the same residue-level epitope in an identical binding pose. Epitope resolution of SPACE2 appears to be similar to that obtained from crystal structures and higher than data from epitope binning methods which struggle to distinguish between antibodies that bind distinct sites but overlap sterically.

SPACE2 outperforms our previous epitope profiling tool SPACE1 (11) and clonotyping when considering the number of antibodies in epitope-consistent multiple-occupancy clusters. Clonotyping is more accurate than SPACE2 but has far lower coverage. The lower accuracy of SPACE2 is explained by the fact that antibodies with similar CDR structures may engage different epitopes if chemical properities of the CDR residues are significantly different.

We also highlight how our methodology allows the detection of functional convergence across populations of antibodies. Across functionally consistent clusters of our largest data set, CoV-AbDab (26), we detect a mean CDRH3 sequence identity as low as 54%. Furthermore, we observe 26 functional clusters containing antibodies from multiple species including human, mouse and rhesus macaque antibodies. In comparison, sequence-based epitope profiling such as clonotyping is severely restricted in grouping sequence diverse antibodies and is not able to link antibodies from different genetic origins and species (9, 10, 45). While it is possible to cluster nanobodies with the SPACE2-HC implementation, when testing on CoV-AbDab we were unable to detect functional convergence to antibodies. No clusters were detected containing both antibodies and nanobodies suggesting the two formats use different binding site structures to engage common epitopes.

SPACE2 clusters antibodies based on the length and structural similarity of all six CDRs. This approach may constrain the detection of functional convergence to some extent as it assumes antibodies require the same length of all six CDRs to engage the same epitope. We tried to address this issue by evaluating two adaptations of SPACE2 that reduce the number of CDRs required to have the same length. An implementation clustering antibodies based on heavy chain structural similarity (SPACE2-HC) caused a drastic decrease in clustering accuracy. This indicates that light chain structures are important for determining antibody binding specificity which is in line with previous findings on the functional selection of light chains (46) and their structural importance (47). Similarly, combining SPACE2 with information from paratope prediction (SPACE2-Paratope) (39) and computing structural similarity only across CDRs predicted to contain paratope residues currently led to fewer functionally consistent clusters.

The ability to detect functional convergence of antibodies will provide valuable insights into the humoral immune response. SPACE2 is able to provide more complete information on public epitopes targeted by antibodies originating from different individuals and species. Whereas previous studies show several public epitopes are largely distinct between species (44), here we identify a number of inter-species clusters. Structural clustering of larger data sets of antibodies isolated from various species will further improve our understanding of differences in their immune responses.

While SPACE2 is computationally more expensive than sequence-based epitope profiling it is tractable for data sets of 10^4^ antibodies, a typical number of sequences obtained from methods such as 10X sequencing (Fig. S4). The rate limiting step of SPACE2 is currently the prediction of antibody structures. Improvements in the speed of structure prediction tools as well as the release of antibody databases containing pre-modelled structures (21) will contribute to reducing the computational cost of structure-based epitope profiling.

Overall, SPACE2 efficiently detects functional convergence of antibodies with highly diverse sequences, genetic lineage and species origins, further illustrating that predicted structures should be considered when investigating the function of antibodies. SPACE2 is openly available on Github (https://github.com/oxpig/SPACE2).

## Author Contributions

FS and CD contributed to conception and design of the study. FS and BA wrote the code for the SPACE2 method. FS performed the data analysis and wrote the manuscript. MR, WW, GG and CD contributed to critical revision of the manuscript.

## Supplementary Materials

**Fig. S1.**
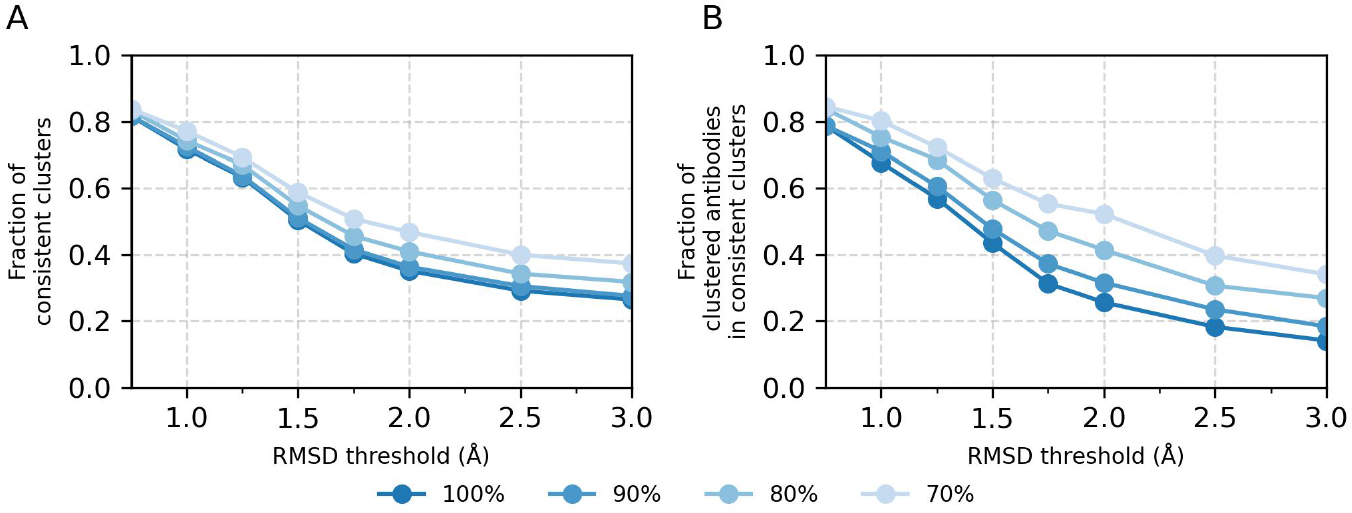
Extent of cluster inconsistency. Accuracy metrics used in this study only capture clusters that exclusively contain antibodies against the same epitope (100% consistency). The two accuracy metrics, (A) the fraction of epitope-consistent clusters and (B) the fraction of clustered antibodies in epitope-consistent clusters, were recalculated considering clusters where at least 90%, 80% and 70% of the members engage the same epitope to be epitope-consistent. This provides a measure of the extent of inconsistency in clusters not captured by the standard performance metrics. A large number of clusters are observed where most antibodies engage the same epitope, but contain a few incorrect antibodies.

**Fig. S2.**
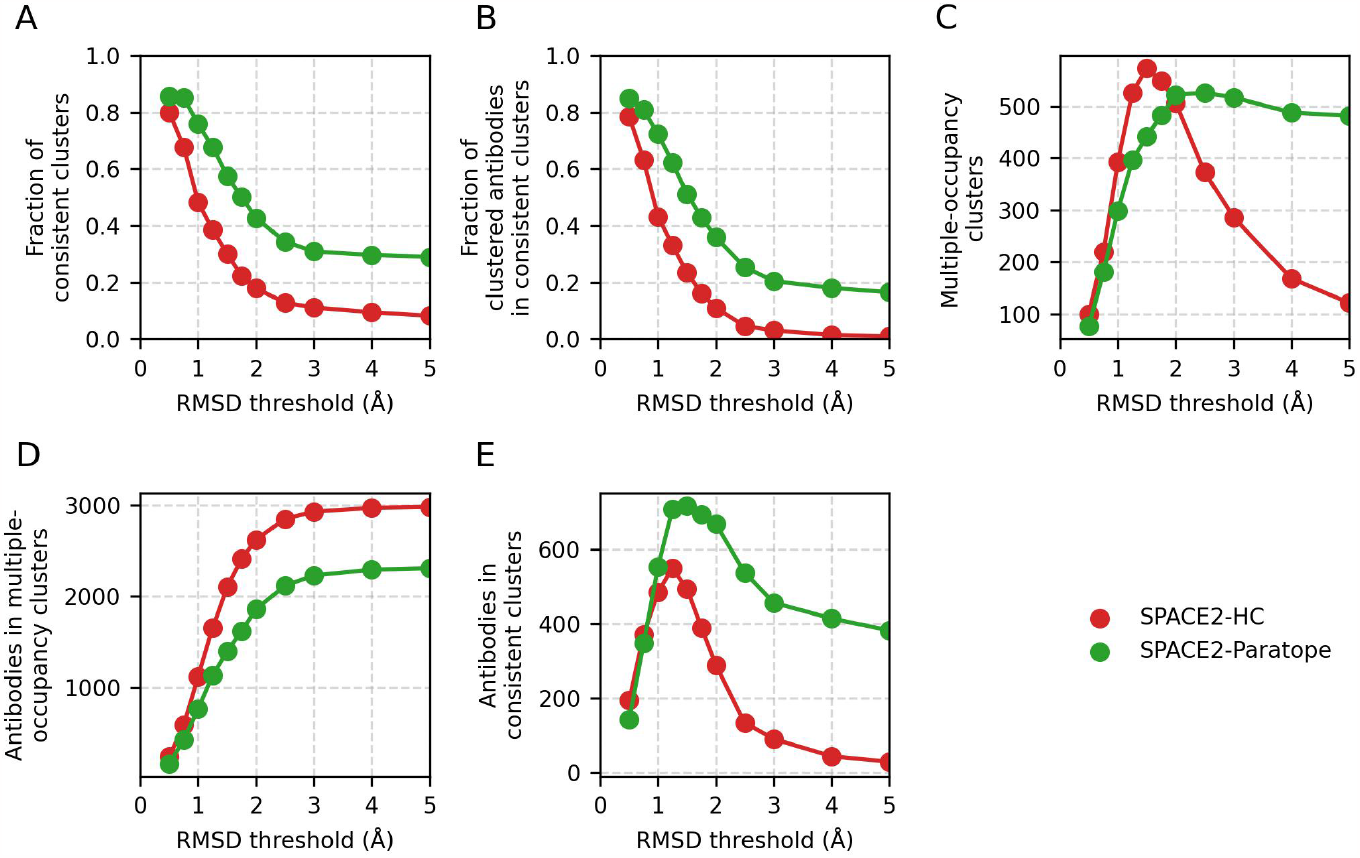
Parameter optimisation of SPACE2 variations. Versions of the algorithm clustering antibodies based on the similarity of heavy chain CDRs (SPACE2-HC) and the similarity of CDRs predicted to contain paratope residues (SPACE2-Paratope) were implemented. As the standard SPACE2 method, both SPACE2-HC and SPACE2-Paratope cluster antibodies using an agglomerative clustering algorithm with a ‘complete’ linkage criterion. A scan of the RMSD threshold parameter was performed to find optimal values. Optimisation was carried out on the Cao et al. (25) training set. The values of the five performance metrics are plotted against evaluated threshold values: (A) fraction of epitope-consistent clusters, (B) fraction of clustered antibodies in epitope-consistent clusters, (C) number of multiple-occupancy clusters, (D) number of antibodies in multiple-occupancy clusters, (E) number of antibodies in epitope-consistent multiple-occupancy clusters. The best clustering results are achieved at a threshold value of 1.25 Å for SPACE2-HC and 1.5 Å for SPACE2-Paratope as defined by the maximum number of antibodies in epitope-consistent clusters.

**Fig. S3.**
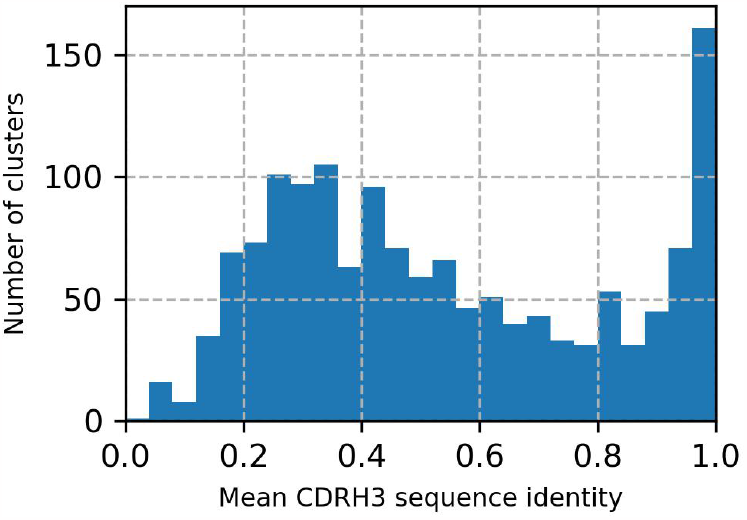
Histogram of the mean CDRH3 sequence identity within domain-consistent SPACE2 clusters from CoV-AbDab. The distribution shows two peaks. The peak close to 1.0 indicates a group of clusters exclusively containing antibodies with almost identical sequences. A second peak at 0.3 indicates the presence of a large number of clusters containing antibodies that are highly diverse in sequence.

**Fig. S4.**
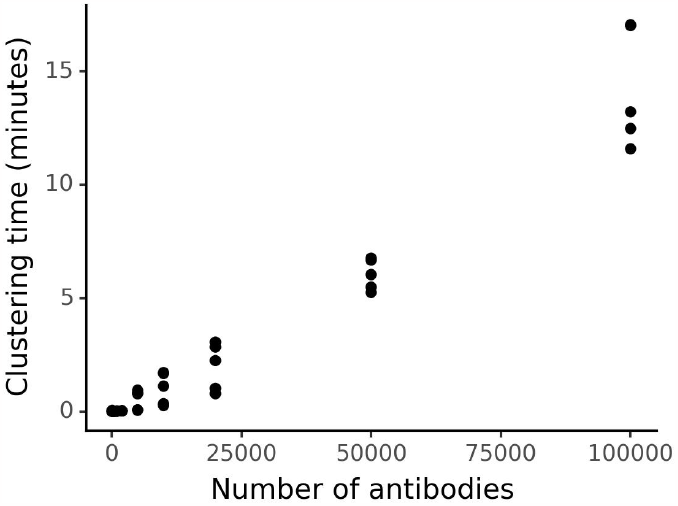
SPACE2 clustering speed. The SPACE2 algorithm functions in four steps: structural modelling of antibodies, grouping by CDR length, computation of the RMSD matrices and agglomerative clustering. Structural modelling with ABodyBuilder2 is currently the rate limiting step, which take around 5 seconds per structure using a Tesla P100 GPU (20). To determine the computational cost of the remaining three steps, we clustered a database of 148,000 paired natural antibody sequences from the Observed Antibody Space database pre-modelled with ABodyBuilder2 (20, 48). The graph shows the time required to run the remaining three steps of the clustering code parallelised over 12 CPUs plotted against the number of antibodies to be clustered. These steps scale roughly at O(n^1.5^) with the number of antibodies (n). It is possible to cluster a data set of 10,000 antibodies with SPACE2 within half a day (including all steps of the algorithm). In comparison, clonotyping has a lower computational cost. Our in-house script for Fv-clonotyping takes approximately 2 seconds to cluster 10,000 antibodies on a single CPU.

**Table S1.** Clustering algorithms and parameter ranges/values evaluated during optimisation.

| Algorithm | Parameter | Range/values | Optimal value |
| --- | --- | --- | --- |
| Greedy clustering | RMSD threshold | 0.5 - 10 Å | 1.25 Å |
| Agglomerative clustering | RMSD threshold | 0.5 - 10 Å | 1.25 Å |
|  | linkage criterion | complete, average, single | complete |
| Affinity propagation | Preferences | -5 - 4 | Median(RMSD matrix) |
| DBSCAN | RMSD threshold | 0.5 - 5 Å | 1 Å |
|  | minimum samples | 2, 5 | 2 |
| OPTICS-xi | RMSD threshold | 0.5 - 5 Å | 1.5 Å |
|  | minimum samples | 2 | 2 |
|  | xi | 0.005 - 0.5 | ≤0.01 |
| OPTICS-DBSCAN | RMSD threshold | 0.5 - 2 Å | 1 Å |
|  | minimum samples | 2 | 2 |
| K-means | Initialisation | random, K-means++ | K-means++ |
| Butina clustering | RMSD threshold | 0.5 - 5 Å | 1 Å |
|  | reordering | True, False | False |

**Table S2.**
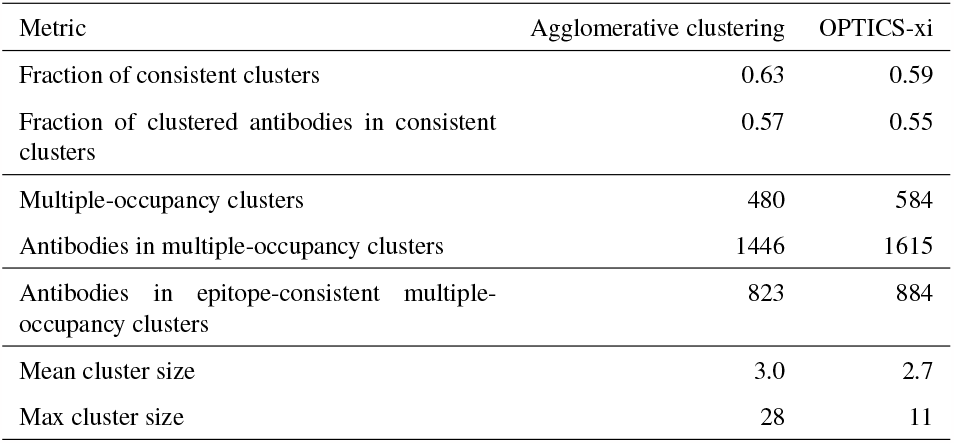
Performance of agglomerative and OPTICS-xi clustering on train set.

| Metric | Agglomerative clustering | OPTICS-xi |
| --- | --- | --- |
| Fraction of consistent clusters | 0.63 | 0.59 |
| Fraction of clustered antibodies in consistent clusters | 0.57 | 0.55 |
| Multiple-occupancy clusters | 480 | 584 |
| Antibodies in multiple-occupancy clusters | 1446 | 1615 |
| Antibodies in epitope-consistent multiple-occupancy clusters | 823 | 884 |
| Mean cluster size | 3.0 | 2.7 |
| Max cluster size | 28 | 11 |

**Table S3.** Comparison of standard SPACE2, SPACE2-HC and SPACE2-Paratope performance metrics on the Cao et al. (25) training set. The best results for each metric are highlighted in bold.

| Metric | Standard SPACE2 | SPACE2-HC | SPACE2-Paratope |
| --- | --- | --- | --- |
| Fraction of consistent clusters | **0.63** | 0.36 | 0.57 |
| Fraction of clustered antibodies in consistent clusters | **0.57** | 0.33 | 0.51 |
| Multiple-occupancy clusters | 480 | **526** | 442 |
| Antibodies in multiple-occupancy clusters | 1446 | **1655** | 1339 |
| Antibodies in epitope-consistent multiple-occupancy clusters | **823** | 550 | 717 |

**Table S4.** Comparison of SPACE1 evaluated at different RMSD thresholds on the Cao et al. (25) training set. The lower threshold leads to better metrics of clustering accuracy. However, a larger threshold improves data coverage and the key metric of antibodies in epitope-consistent multiple-occupancy clusters. The best results achieved by SPACE1 for each metric are highlighted in bold. Values achieved by SPACE2 are shown for comparison.

| Metric | SPACE1 0.75 Å | SPACE1 1.25 Å | SPACE2 |
| --- | --- | --- | --- |
| Fraction of consistent clusters | **0.75** | 0.64 | 0.63 |
| Fraction of clustered antibodies in consistent clusters | **0.68** | 0.58 | 0.57 |
| Multiple-occupancy clusters | 251 | **314** | 480 |
| Antibodies in multiple-occupancy clusters | 723 | **935** | 1446 |
| Antibodies in epitope-consistent multiple-occupancy clusters | 494 | **538** | 823 |
| Fraction of clusters containing >1 VH-clonotypes | 0.55 | **0.58** | 0.55 |
| Mean CDRH3 sequence identity | 0.70 | **0.67** | 0.66 |

## Bibliography

1. Konstantinos Tsioris, Namita T. Gupta, Adebola O. Ogunniyi, Ross M. Zimnisky, Feng Qian, Yi Yao, Xiaomei Wang, Joel N. H. Stern, Raj Chari, Adrian W. Briggs, Christopher R. Clouser, Francois Vigneault, George M. Church, Melissa N. Garcia, Kristy O. Murray, Ruth R. Montgomery, Steven H. Kleinstein, and J. Christopher Love. Neutralizing antibodies against West Nile virus identified directly from human B cells by single-cell analysis and next generation sequencing. Integrative Biology, 7(12):1587–1597, 2015. ISSN 1757-9694, 1757-9708. doi: 10.1039/C5IB00169B.

2. R. J. M. Bashford-Rogers, L. Bergamaschi, E. F. McKinney, D. C. Pombal, F. Mescia, J. C. Lee, D. C. Thomas, S. M. Flint, P. Kellam, D. R. W. Jayne, P. A. Lyons, and K. G. C. Smith. Analysis of the B cell receptor repertoire in six immune-mediated diseases. Nature, 574(7776):122–126, October 2019. ISSN 1476-4687. doi: 10.1038/s41586-019-1595-3.

3. Bryan Briney, Anne Inderbitzin, Collin Joyce, and Dennis R. Burton. Commonality despite exceptional diversity in the baseline human antibody repertoire. Nature, 566(7744):393–397, February 2019. ISSN 1476-4687. doi: 10.1038/s41586-019-0879-y.

4. Sai T. Reddy, Xin Ge, Aleksandr E. Miklos, Randall A. Hughes, Seung Hyun Kang, Kam Hon Hoi, Constantine Chrysostomou, Scott P. Hunicke-Smith, Brent L. Iverson, Philip W. Tucker, Andrew D. Ellington, and George Georgiou. Monoclonal antibodies isolated without screening by analyzing the variable-gene repertoire of plasma cells. Nature Biotechnology, 28(9):965–969, September 2010. ISSN 1546-1696. doi: 10.1038/nbt.1673.

5. Jiang Zhu, Gilad Ofek, Yongping Yang, Baoshan Zhang, Mark K. Louder, Gabriel Lu, Krisha McKee, Marie Pancera, Jeff Skinner, Zhenhai Zhang, Robert Parks, Joshua Eudailey, Krissey E. Lloyd, Julie Blinn, S. Munir Alam, Barton F. Haynes, Melissa Simek, Dennis R. Burton, Wayne C. Koff, NISC Comparative Sequencing Program, James C. Mullikin, John R. Mascola, Lawrence Shapiro, Peter D. Kwong, Jesse Becker, Betty Benjamin, Robert Blakesley, Gerry Bouffard, Shelise Brooks, Holly Coleman, Mila Dekhtyar, Michael Gregory, Xiaobin Guan, Jyoti Gupta, Joel Han, April Hargrove, Shi-ling Ho, Taccara Johnson, Richelle Legaspi, Sean Lovett, Quino Maduro, Cathy Masiello, Baishali Maskeri, Jenny McDowell, Casandra Montemayor, James Mullikin, Morgan Park, Nancy Riebow, Karen Schandler, Brian Schmidt, Christina Sison, Mal Stantripop, James Thomas, Pam Thomas, Meg Vemulapalli, and Alice Young. Mining the antibodyome for HIV-1–neutralizing antibodies with next-generation sequencing and phylogenetic pairing of heavy/light chains. Proceedings of the National Academy of Sciences, 110(16):6470–6475, April 2013. ISSN 0027-8424, 1091-6490. doi: 10.1073/pnas.1219320110.

6. Yi-Chun Hsiao, Yonglei Shang, Danielle M. DiCara, Angie Yee, Joyce Lai, Si Hyun Kim, Diego Ellerman, Racquel Corpuz, Yongmei Chen, Sharmila Rajan, Hao Cai, Yan Wu, Dhaya Seshasayee, and Isidro Hötzel. Immune repertoire mining for rapid affinity optimization of mouse monoclonal antibodies. mAbs, 11(4):735–746, May 2019. ISSN 1942-0862. doi: 10.1080/19420862.2019.1584517.

7. Johan Nilvebrant and Johan Rockberg. An Introduction to Epitope Mapping. In Johan Rockberg and Johan Nilvebrant, ditors, Epitope Mapping Protocols, Methods in Molecular Biology, pages 1–10. Springer, New York, NY, 2018. ISBN 9781493978410. doi: 10.1007/978-1-4939-7841-0_1.

8. Yasmina N. Abdiche, Dan S. Malashock, Alanna Pinkerton, and Jaume Pons. Exploring blocking assays using Octet, ProteOn, and Biacore biosensors. Analytical Biochemistry, 386(2):172–180, March 2009. ISSN 0003-2697. doi: 10.1016/j.ab.2008.11.038.

9. Laura López-Santibáñez-Jácome, S. Eréndira Avendaño-Vázquez, and Carlos Fabián Flores-Jasso. The Pipeline Repertoire for Ig-Seq Analysis. Frontiers in Immunology, 10:899, April 2019. ISSN 1664-3224. doi: 10.3389/fimmu.2019.00899.

10. Victor Greiff, Enkelejda Miho, Ulrike Menzel, and Sai T. Reddy. Bioinformatic and Statistical Analysis of Adaptive Immune Repertoires. Trends in Immunology, 36(11):738–749, November 2015. ISSN 14714906. doi: 10.1016/j.it.2015.09.006.

11. Sarah A. Robinson, Matthew I. J. Raybould, Constantin Schneider, Wing Ki Wong, Claire Marks, and Charlotte M. Deane. Epitope profiling using computational structural modelling demonstrated on coronavirus-binding antibodies. PLOS Computational Biology, 17 (12):e1009675, December 2021. ISSN 1553-7358. doi: 10.1371/journal.pcbi.1009675.

12. Wing Ki Wong, Sarah A. Robinson, Alexander Bujotzek, Guy Georges, Alan P. Lewis, Jiye Shi, James Snowden, Bruck Tad-dese, and Charlotte M. Deane. Ab-Ligity: identifying sequence-dissimilar antibodies that bind to the same epitope. mAbs, 13 (1):1873478, January 2021. ISSN 1942-0862, 1942-0870. doi: 10.1080/19420862.2021.1873478.

13. Johannes F. Scheid, Hugo Mouquet, Beatrix Ueberheide, Ron Diskin, Florian Klein, Thiago Y. K. Oliveira, John Pietzsch, David Fenyo, Alexander Abadir, Klara Velinzon, Arlene Hurley, Sunnie Myung, Farid Boulad, Pascal Poignard, Dennis R. Burton, Florencia Pereyra, David D. Ho, Bruce D. Walker, Michael S. Seaman, Pamela J. Bjorkman, Brian T. Chait, and Michel C. Nussen-zweig. Sequence and Structural Convergence of Broad and Potent HIV Antibodies That Mimic CD4 Binding. Science, 333(6049): 1633–1637, September 2011. ISSN 0036-8075, 1095-9203. doi: 10.1126/science.1207227.

14. Pramila Rijal, Sean C. Elias, Samara Rosendo Machado, Julie Xiao, Lisa Schimanski, Victoria O’Dowd, Terry Baker, Emily Barry, Simon C. Mendelsohn, Catherine J. Cherry, Jing Jin, Geneviève M. Labbé, Francesca R. Donnellan, Tommy Rampling, Stuart Dowall, Emma Rayner, Stephen Findlay-Wilson, Miles Carroll, Jia Guo, Xiao-Ning Xu, Kuan-Ying A. Huang, Ayato Takada, Gillian Burgess, David McMillan, Andy Popplewell, Daniel J. Lightwood, Simon J. Draper, and Alain R. Townsend. Therapeutic Monoclonal Antibodies for Ebola Virus Infection Derived from Vaccinated Humans. Cell Reports, 27(1):172–186.e7, April 2019. ISSN 2211-1247. doi: 10.1016/j.celrep.2019.03.020.

15. M. Gordon Joyce, Adam K. Wheatley, Paul V. Thomas, Gwo-Yu Chuang, Cinque Soto, Robert T. Bailer, Aliaksandr Druz, Ivelin S. Georgiev, Rebecca A. Gillespie, Masaru Kanekiyo, Wing-Pui Kong, Kwanyee Leung, Sandeep N. Narpala, Madhu S. Prabhakaran, Eun Sung Yang, Baoshan Zhang, Yi Zhang, Mangaiarkarasi Asokan, Jeffrey C. Boyington, Tatsiana Bylund, Sam Darko, Christopher R. Lees, Amy Ransier, Chen-Hsiang Shen, Lingshu Wang, James R. Whittle, Xueling Wu, Hadi M. Yassine, Celia Santos, Yumiko Matsuoka, Yaroslav Tsybovsky, Ulrich Baxa, James C. Mullikin, Kanta Subbarao, Daniel C. Douek, Barney S. Graham, Richard A. Koup, Julie E. Ledgerwood, Mario Roederer, Lawrence Shapiro, Peter D. Kwong, John R. Mascola, and Adrian B. McDermott. Vaccine-Induced Antibodies that Neutralize Group 1 and Group 2 Influenza A Viruses. Cell, 166(3):609–623, July 2016. ISSN 0092-8674. doi: 10.1016/j.cell.2016.06.043.

16. Eve Richardson, Jacob D. Galson, Paul Kellam, Dominic F. Kelly, Sarah E. Smith, Anne Palser, Simon Watson, and Charlotte M. Deane. A computational method for immune repertoire mining that identifies novel binders from different clonotypes, demonstrated by identifying anti-pertussis toxoid antibodies. mAbs, 13 (1):1869406, January 2021. ISSN 1942-0862, 1942-0870. doi: 10.1080/19420862.2020.1869406.

17. John Jumper, Richard Evans, Alexander Pritzel, Tim Green, Michael Figurnov, Olaf Ronneberger, Kathryn Tunyasuvunakool, Russ Bates, Augustin Žídek, Anna Potapenko, Alex Bridgland, Clemens Meyer, Simon A. A. Kohl, Andrew J. Ballard, Andrew Cowie, Bernardino Romera-Paredes, Stanislav Nikolov, Rishub Jain, Jonas Adler, Trevor Back, Stig Petersen, David Reiman, Ellen Clancy, Michal Zielinski, Martin Steinegger, Michalina Pacholska, Tamas Berghammer, Sebastian Bodenstein, David Silver, Oriol Vinyals, Andrew W. Senior, Koray Kavukcuoglu, Pushmeet Kohli, and Demis Hassabis. Highly accurate protein structure prediction with AlphaFold. Nature, 596(7873):583–589, August 2021. ISSN 1476-4687. doi: 10.1038/s41586-021-03819-2.

18. Minkyung Baek, Frank DiMaio, Ivan Anishchenko, Justas Dauparas, Sergey Ovchinnikov, Gyu Rie Lee, Jue Wang, Qian Cong, Lisa N. Kinch, R. Dustin Schaeffer, Claudia Millán, Hahnbeom Park, Carson Adams, Caleb R. Glassman, Andy DeGiovanni, Jose H. Pereira, Andria V. Rodrigues, Alberdina A. van Dijk, Ana C. Ebrecht, Diederik J. Opperman, Theo Sagmeister, Christoph Buhlheller, Tea Pavkov-Keller, Manoj K. Rathinaswamy, Udit Dalwadi, Calvin K. Yip, John E. Burke, K. Christopher Garcia, Nick V. Grishin, Paul D. Adams, Randy J. Read, and David Baker. Accurate prediction of protein structures and interactions using a three-track neural network. Science, 373(6557):871–876, August 2021. ISSN 0036-8075, 1095-9203. doi: 10.1126/science.abj8754.

19. Zeming Lin, Halil Akin, Roshan Rao, Brian Hie, Zhongkai Zhu, Wenting Lu, Allan dos Santos Costa, Maryam Fazel-Zarandi, Tom Sercu, Sal Candido, and Alexander Rives. Language models of protein sequences at the scale of evolution enable accurate structure prediction, July 2022.

20. Brennan Abanades, Wing Ki Wong, Fergus Boyles, Guy Georges, Alexander Bujotzek, and Charlotte M. Deane. ImmuneBuilder: Deep-Learning models for predicting the structures of immune proteins, December 2022.

21. Brennan Abanades, Guy Georges, Alexander Bujotzek, and Char-lotte M Deane. ABlooper: fast accurate antibody CDR loop structure prediction with accuracy estimation. Bioinformatics, 38(7): 1877–1880, March 2022. ISSN 1367-4803, 1460-2059. doi: 10.1093/bioinformatics/btac016.

22. Jeffrey A. Ruffolo, Carlos Guerra, Sai Pooja Mahajan, Jeremias Sulam, and Jeffrey J. Gray. Geometric potentials from deep learning improve prediction of CDR H3 loop structures. Bioinformatics (Oxford, England), 36(Suppl_1):i268–i275, July 2020. ISSN 1367-4811. doi: 10.1093/bioinformatics/btaa457.

23. Jeffrey A. Ruffolo, Jeremias Sulam, and Jeffrey J. Gray. Antibody structure prediction using interpretable deep learning. Patterns, 3 (2):100406, February 2022. ISSN 26663899. doi: 10.1016/j.patter.2021.100406.

24. Jeffrey A. Ruffolo, Lee-Shin Chu, Sai Pooja Mahajan, and Jeffrey J. Gray. Fast, accurate antibody structure prediction from deep learning on massive set of natural antibodies, April 2022.

25. Yunlong Cao, Fanchong Jian, Jing Wang, Yuanling Yu, Weiliang Song, Ayijiang Yisimayi, Jing Wang, Ran An, Xiaosu Chen, Na Zhang, Yao Wang, Peng Wang, Lijuan Zhao, Haiyan Sun, Lingling Yu, Sijie Yang, Xiao Niu, Tianhe Xiao, Qingqing Gu, Fei Shao, Xiaohua Hao, Yanli Xu, Ronghua Jin, Zhongyang Shen, Youchun Wang, and Xiaoliang Sunney Xie. Imprinted SARS-CoV-2 humoral immunity induces convergent Omicron RBD evolution. Nature, 614(7948):521–529, February 2023. ISSN 1476-4687. doi: 10.1038/s41586-022-05644-7.

26. Matthew I J Raybould, Aleksandr Kovaltsuk, Claire Marks, and Charlotte M Deane. CoV-AbDab: the coronavirus antibody database. Bioinformatics, 37(5):734–735, May 2021. ISSN 1367-4803, 1460-2059. doi: 10.1093/bioinformatics/btaa739.

27. Constantin Schneider, Matthew I J Raybould, and Charlotte M Deane. SAbDab in the age of biotherapeutics: updates including SAbDab-nano, the nanobody structure tracker. Nucleic Acids Research,50(D1):D1368–D1372, January 2022. ISSN 0305-1048, 1362-4962. doi: 10.1093/nar/gkab1050.

28. James Dunbar, Konrad Krawczyk, Jinwoo Leem, Terry Baker, Angelika Fuchs, Guy Georges, Jiye Shi, and Charlotte M. Deane. SAbDab: the structural antibody database. Nucleic Acids Research, 42(D1):D1140–D1146, January 2014. ISSN 0305-1048, 1362-4962. doi: 10.1093/nar/gkt1043.

29. Jinwoo Leem, James Dunbar, Guy Georges, Jiye Shi, and Charlotte M. Deane. ABodyBuilder: Automated antibody structure prediction with data-driven accuracy estimation. mAbs, 8(7):1259–1268, October 2016. ISSN 1942-0870. doi: 10.1080/19420862.2016.1205773.

30. Yoonjoo Choi and Charlotte M. Deane. FREAD revisited: Accurate loop structure prediction using a database search algorithm: Loop Structure Prediction Using a Database Search. Proteins: Structure, Function, and Bioinformatics, 78(6):1431–1440, May 2010. ISSN 08873585. doi: 10.1002/prot.22658.

31. Fionn Murtagh and Pedro Contreras. Algorithms for hierarchical clustering: an overview. WIREs Data Mining and Knowledge Discovery, 2(1):86–97, January 2012. ISSN 1942-4787, 1942-4795. doi: 10.1002/widm.53.

32. Brendan J. Frey and Delbert Dueck. Clustering by Passing Messages Between Data Points. Science, 315(5814):972–976, February 2007. ISSN 0036-8075, 1095-9203. doi: 10.1126/science.1136800.

33. Erich Schubert, Jörg Sander, Martin Ester, Hans Peter Kriegel, and Xiaowei Xu. DBSCAN Revisited, Revisited: Why and How You Should (Still) Use DBSCAN. ACM Transactions on Database Systems, 42(3):19:1–19:21, July 2017. ISSN 0362-5915. doi: 10.1145/3068335.

34. Mihael Ankerst, Markus M. Breunig, Hans-Peter Kriegel, and Jörg Sander. OPTICS: ordering points to identify the clustering structure. ACM SIGMOD Record, 28(2):49–60, June 1999. ISSN 0163-5808. doi: 10.1145/304181.304187.

35. J. MacQueen. Some methods for classification and analysis of multivariate observations. Proceedings of the Fifth Berkeley Symposium on Mathematical Statistics and Probability, Volume 1: Statistics, 5.1:281–298, January 1967.

36. Fabian Pedregosa, Gaël Varoquaux, Alexandre Gramfort, Vincent Michel, Bertrand Thirion, Olivier Grisel, Mathieu Blondel, Peter Prettenhofer, Ron Weiss, Vincent Dubourg, Jake Vanderplas, Alexandre Passos, David Cournapeau, Matthieu Brucher, Matthieu Perrot, and Édouard Duchesnay. Scikit-learn: Machine Learning in Python. Journal of Machine Learning Research, 12(85):2825–2830, 2011. ISSN 1533-7928.

37. Darko Butina. Unsupervised Data Base Clustering Based on Day-light’s Fingerprint and Tanimoto Similarity: A Fast and Automated Way To Cluster Small and Large Data Sets. Journal of Chemical Information and Computer Sciences, 39(4):747–750, July 1999. ISSN 0095-2338. doi: 10.1021/ci9803381.

38. Greg Landrum. RDKit: Open-source cheminformatics, 2006.

39. Lewis Chinery, Newton Wahome, Iain Moal, and Charlotte M Deane. Paragraph—antibody paratope prediction using graph neural networks with minimal feature vectors. Bioinformatics, 39(1):btac732, January 2023. ISSN 1367-4811. doi: 10.1093/bioinformatics/btac732.

40. Marie-Paule Lefranc, Christelle Pommié, Manuel Ruiz, Véronique Giudicelli, Elodie Foulquier, Lisa Truong, Valérie Thouvenin-Contet, and Gérard Lefranc. IMGT unique numbering for immunoglobulin and T cell receptor variable domains and Ig superfamily V-like domains. Developmental & Comparative Immunology, 27(1):55–77, January 2003. ISSN 0145-305X. doi: 10.1016/S0145-305X(02)00039-3.

41. Benjamin North, Andreas Lehmann, and Roland L. Dunbrack. A New Clustering of Antibody CDR Loop Conformations. Journal of Molecular Biology, 406(2):228–256, February 2011. ISSN 00222836. doi: 10.1016/j.jmb.2010.10.030.

42. Jeffrey E. Lee, Marnie L. Fusco, Ann J. Hessell, Wendelien B. Oswald, Dennis R. Burton, and Erica Ollmann Saphire. Structure of the Ebola virus glycoprotein bound to an antibody from a human survivor. Nature, 454(7201):177–182, July 2008. ISSN 1476-4687. doi: 10.1038/nature07082.

43. Ammar Tareen and Justin B Kinney. Logomaker: beautiful sequence logos in Python. Bioinformatics, 36(7):2272–2274, April 2020. ISSN 1367-4803, 1460-2059. doi: 10.1093/bioinformatics/btz921.

44. Ellen L. Shrock, Richard T. Timms, Tomasz Kula, Elijah L. Mena, Anthony P. West, Rui Guo, I-Hsiu Lee, Alexander A. Cohen, Lindsay G. A. McKay, Caihong Bi, Yumei Leng, Eric Fujimura, Felix Horns, Mamie Li, Duane R. Wesemann, Anthony Griffiths, Benjamin E. Gewurz, Pamela J. Bjorkman, and Stephen J. Elledge. Germline-encoded amino acid–binding motifs drive immunodominant public antibody responses. Science, 380(6640):eadc9498, April 2023. ISSN 0036-8075, 1095-9203. doi: 10.1126/science.adc9498.

45. Matthew I. J. Raybould, Anthony R. Rees, and Charlotte M. Deane. Current strategies for detecting functional convergence across B-cell receptor repertoires. mAbs, 13(1):1996732, January 2021. ISSN 1942-0862. doi: 10.1080/19420862.2021.1996732.

46. David B. Jaffe, Payam Shahi, Bruce A. Adams, Ashley M. Chrisman, Peter M. Finnegan, Nandhini Raman, Ariel E. Royall, FuNien Tsai, Thomas Vollbrecht, Daniel S. Reyes, N. Lance Hepler, and Wyatt J. McDonnell. Functional antibodies exhibit light chain coherence. Nature, 611(7935):352–357, November 2022. ISSN 1476-4687. doi: 10.1038/s41586-022-05371-z.

47. Bora Guloglu and Charlotte M. Deane. Specific attributes of the VL domain influence both the structure and structural variability of CDR-H3 through steric effects, May 2023.

48. Tobias H. Olsen, Fergus Boyles, and Charlotte M. Deane. Observed Antibody Space: A diverse database of cleaned, annotated, and translated unpaired and paired antibody sequences. Protein Science, 31(1):141–146, January 2022. ISSN 0961-8368, 1469-896X. doi: 10.1002/pro.4205.

